# Signalome-wide mapping of the NFκB pathway in T-cells reveals novel targets for immunotherapy

**DOI:** 10.64898/2026.02.16.706160

**Authors:** Joseph Clarke, Haoqi Chen, Cristina Tormo Garcia, Ewa Basiarz, Mateusz Kotowski, Ana Mafalda Santos, Mai T Voung, Paige Sherman, Jia Xin Li, Christopher J Tape, Simon J Davis, Evangelia Petsalaki, Sumana Sharma

## Abstract

Cell signalling networks govern fundamental cellular processes yet remain incompletely defined. Moreover, what is known is biased toward a limited subset of well-characterised components. Phosphoprotein-based interrogation methods, including mass spectrometry and targeted phosphosite panels, have limited utility in physiological settings dependent on cell–cell interactions because the signalling fluxes can be difficult to detect despite producing robust functional responses. Here we developed a perturbation-based experimental framework that infers signalling pathway architecture using quantitative functional outputs rather than direct measurements of effector state, e.g., phosphorylation levels. Using antigen-specific, NF-κB–GFP reporter-expressing transformed T-cells co-cultured with cellular targets, we performed an arrayed CRISPR–Cas9 screen targeting a curated signalome of kinases, phosphatases, adaptor and scaffolding proteins, totalling 706 genes. Quantitative effect-size profiling recovered canonical T-cell receptor regulators and revealed unequal, family-specific patterns of control over NF-κB activation. Comparing T-cell stimulation with low- and high-affinity antigen uncovered signal-strength-dependent buffering of proximal signalling nodes, exemplified by reduced sensitivity to perturbation of *LCK* under high-intensity stimulation. Targeted perturbation in primary human CD8⁺ T-cells validated our findings and identified *TRRAP* and *CTDSPL2* as negative regulators of T-cell effector output, whose disruption enhanced cytotoxicity, degranulation, and cytokine production in both polyclonal and TCR-engineered T cells. Together, these results establish a scalable strategy for mapping signalling pathway architecture in the setting of physiological T-cell activation.

## Introduction

Cellular signalling is crucial for fundamental processes such as growth, proliferation, differentiation, survival, metabolism, and specialised effector responses. Although powerful technologies such as CyTOF, LC–MS-based phosphoproteomics, and targeted phospho-panels have significantly advanced our ability to interrogate signalling events, our understanding of cellular signalling network structure and regulation remains incomplete. Moreover, current knowledge is disproportionately focused on a limited subset of well-studied components, leaving large portions of the signalling landscape poorly characterised.(*1*). This bias is particularly pronounced for kinases and phosphatases, the principal enzymatic drivers of signal transduction, for which an estimated 30-50% of cellular targets remain unknown, indicating that large regions of the signalling landscape remain unexplored (*2*).

Most contemporary approaches to studying signalling rely on direct measurement of molecular intermediates, most commonly phosphorylation events detected by antibody-based assays or mass spectrometry-based phosphoproteomics. Advances in enrichment-based LC-MS have enabled increasingly global profiling of signalling states (*3–8*), and these approaches have been extended through integration with genetic or pharmacological perturbations to infer causal relationships, exemplified by resources such as the Connectivity Map (CMap) (*9*). Despite these advances, molecular readout-based approaches remain limited in their ability to capture signalling behaviour under physiological conditions. They are biased toward abundant or highly dynamic phosphorylation events, require large numbers of cells, and rely heavily on curated pathway annotations for interpretation, reinforcing existing knowledge rather than revealing new regulatory logic. Consequently, signalling components that exert modest, distributed, or context-dependent effects on pathway output may be systematically overlooked.

These limitations are particularly pronounced in systems where signalling is spatially constrained and dependent on direct cell–cell interactions. Most phosphoproteomic and kinase-activity-based studies are performed on bulk cell lysates, restricting their applicability in such contexts. T-cell activation exemplifies this challenge: apart from a small number of SILAC-based co-culture studies (*10–12*), much of the available signalling data relies on supraphysiological stimulation applied outside native cellular contexts to generate detectable responses (*5*, *13*). Single-cell approaches such as mass cytometry (CyTOF) partially address population averaging (*14–17*). However, these approaches also encounter biological limits. In direct T cell–target cell interactions, single-cell mass cytometric analyses reveal remarkably limited signalling flux, often indistinguishable from unstimulated states, in contrast to pharmacological induced T-cell activation (*18*). Notably, despite these modest detectable fluxes, T cells mount robust functional responses, indicating that phosphoprotein-based measurements fail to capture physiologically relevant signalling dynamics.

In contrast, functional outputs integrate signalling information over time and across molecular layers. This suggests an alternative strategy for interrogating signalling networks: rather than measuring signalling intermediates directly, signalling architecture can be inferred by systematically perturbing signalling components and quantifying their impact on downstream functional outputs.

Here, we implement this concept through a perturbation-based framework that reconstructs signalling dependencies directly from functional readouts. By combining arrayed CRISPR-Cas9-mediated gene disruption with a quantitative NF-κB reporter assay, we systematically interrogate the contribution of kinases, phosphatases, and adaptor proteins to signalling pathway transmission, and subsequent transcription factor activity, in a physiologically relevant, cell–cell interaction context. This approach enables quantitative assessment of graded effects across hundreds of signalling components, revealing how control is distributed across a signalling network rather than identifying only a small number of essential nodes.

Applying this strategy to T-cell receptor signalling, we generate a functional dependency map of NF-κB activation that reveals hierarchical control, distributed engagement across signalling families, and context-dependent buffering of proximal signalling nodes as a function of ligand affinity. Extending these findings to primary human CD8⁺ T cells, we identify previously unrecognized negative regulators of T-cell function, including TRRAP and CTDSPL2, whose disruption enhances cytotoxic activity and effector responses. Together, this work establishes a generalisable framework for reconstructing signalling architecture from functional outputs in physiological contexts.

## Results

A functional 1G4-TCR^+^ Jurkat NF-kB reporter system enables analysis of T-cell receptor signalling under physiological stimulation conditions.

To enable systematic interrogation of T-cell signalling through functional outputs, we generated a Jurkat T-cell reporter line in which NF-κB activity is monitored via GFP expression downstream of antigen-specific T-cell receptor (TCR) engagement. NF-κB was selected as a core transcriptional output of T-cell activation, integrating signals from multiple proximal and distal signalling pathways. Jurkat cells were first engineered to stably express an NF-κB-driven GFP reporter (**Supplementary Fig. 1A-B)**. To ensure antigen-specific activation, endogenous TCR expression was ablated by CRISPR targeting of *TRAC* and *TRBC*, and cells were subsequently transduced with lentivirus encoding the 1G4 TCR (**Supplementary Figure 1C**), which recognizes the cancer-testis antigen NY-ESO-1_157-165_ (henceforth referred to simply as NY-ESO-1) peptide presented by HLA-A*02:01, followed by introduction of constitutive Cas9 expression to facilitate CRISPR-Cas9-mediated genetic perturbation (**Supplementary Fig. 1D**).

This configuration establishes a defined receptor-ligand interaction that mimics physiological T-cell-target cell engagement. When co-cultured with A375 melanoma cells pulsed with NY-ESO-1 peptides, 1G4-TCR⁺ Jurkat reporter cells exhibited robust NF-κB–dependent GFP induction, whereas TCR-deficient parental cells remained unresponsive (**Fig. 1A–C**). Upon the use of NY-ESO-1 altered peptide ligands (6T, 3Y, 3I and 9V) to initiate Jurkat T-cell activation, GFP expression scaled with peptide affinity, demonstrating sensitivity across a physiologically relevant range of TCR stimulation strengths (**Fig. 1B,C**).

**Figure 1:**
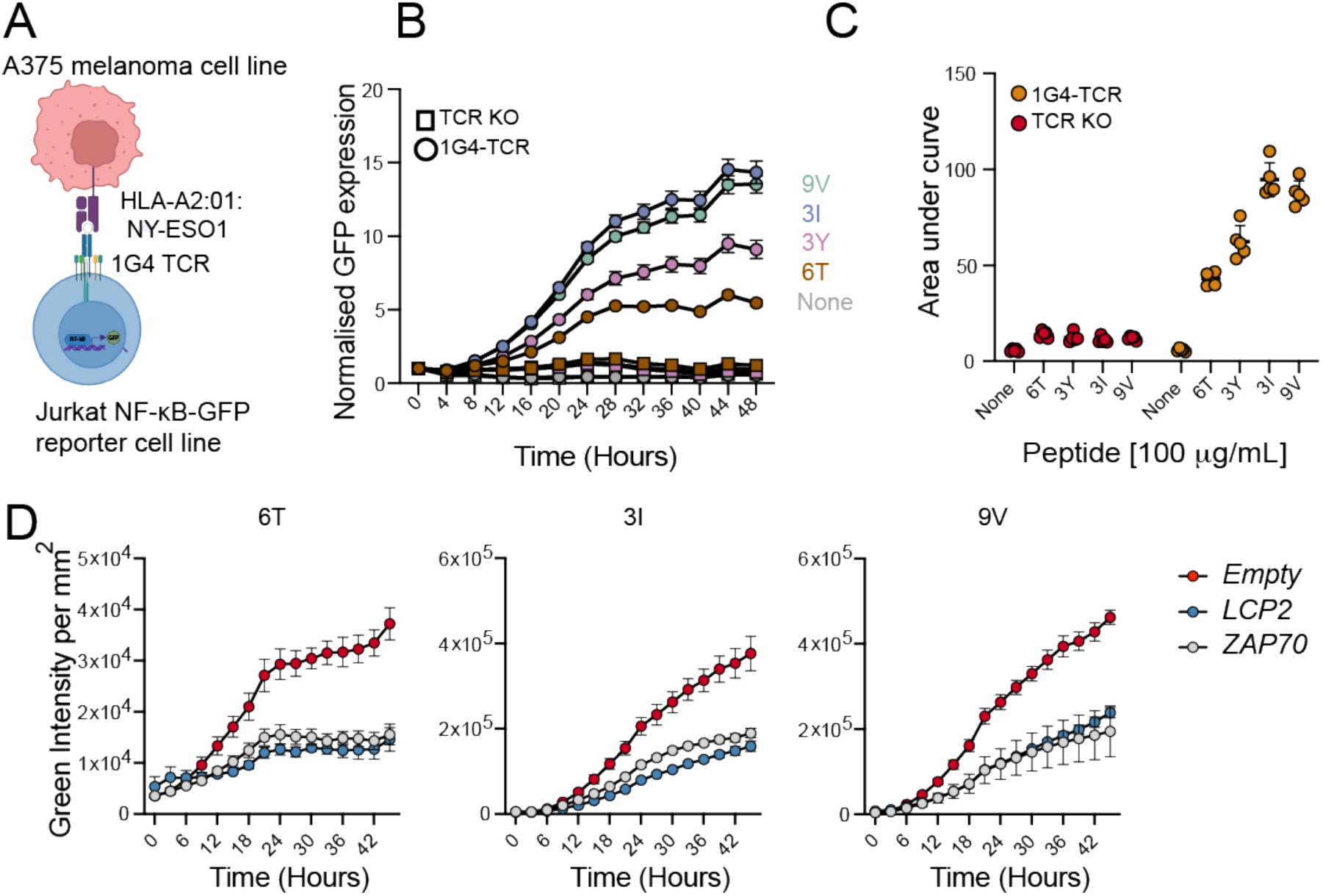
Establishing a peptide:MHC specific cell-cell interaction system to report functional activity of NFkB in T cells A: Schematic representing the co-culture system between Jurkat-NFkB-GFP reporter cell lines expressing the 1G4-TCR, and A375 melanoma cells pulsed with NY-ESO-1 peptides. **B:** Time course IncuCyte measurements of NFkB-GFP intensities from either 1G4-TCR^+^ Jurkat reporter lines (filled circles) or TCR-KO parental lines (filled squares). All NY-ESO-1 altered peptide ligands were used at final concentrations of 100µg/mL. **C:** Area under the curve (AUC) analysis of the data shown in panel B. **D:** Lentiviral sgRNA targeting of key TCR-signalling components *LCP2* or *ZAP70* in 1G4-TCR^+^ Jurkat reporters prevents GFP expression upon co-culture with NY-ESO-1 peptide pulsed A375 melanoma cells. All peptides were used at final concentrations of 100µg/mL, and time course GFP measurements were performed via IncuCyte imaging. All data shown from a minimum of three independent experiments.

To confirm that reporter activation reflected canonical TCR signalling, we genetically disrupted key proximal signalling components. CRISPR-mediated targeting of established TCR signalling mediators, including *LCP2* and *ZAP70*, significantly reduced NF-κB reporter activity upon stimulation with both high- and low-affinity antigen (**Fig. 1D**). These results confirm that NF-κB reporter activation depends on intact TCR-proximal signalling.

Together, these data establish a genetically tractable, antigen-specific Jurkat reporter system that reliably converts physiologically relevant TCR engagement into a quantitative functional readout of NF-κB activation. This platform provides a robust foundation for systematic perturbation-based analysis of signalling dependencies in a cell–cell interaction context, where direct phosphoprotein measurements are known to be challenging.

A signalome-wide CRISPR-Cas9 screen reveals a quantitative map of NF-κB signalling in T cells To enable systematic interrogation of T-cell signalling, we first defined a comprehensive “signalome” comprising kinases, phosphatases, and key adaptor proteins. We combined a curated list of 556 human kinases with 136 protein phosphatases defined by the HUGO Gene Nomenclature Committee (*19*), and filtered this set for genes expressed across multiple immune and cancer cell models using bulk RNA-seq data from Cell Model Passports (TPM > 0.3; Jurkat, MOLT-4, Daudi, THP-1, and two colorectal cancer organoids) (*20*). To further enrich for signalling components relevant to T cells, we included genes annotated in KEGG pathways associated with TCR signalling, JAK-STAT, calcium signalling, actin regulation, phosphatidylinositol signalling, and MAPK signalling (*21*), as well as all genes encoding for SH2 domain-containing proteins, to ensure coverage of adaptor and scaffolding molecules. This yielded a final library of 706 genes, each targeted with two sgRNAs for CRISPR-Cas9-mediated perturbation (**Supplementary Table 1**).

To place this signalome in the context of existing knowledge, we integrated transcriptional and perturbation-based datasets from diverse T-cell contexts. Analysis of published expression datasets spanning multiple activation timepoints in CD8⁺ and pan-T cells (*22–25*) revealed that 446 signalome genes are dynamically regulated at the transcriptional level (**Fig. 2A; Supplementary Table 2**). In parallel, we mapped hits from published genome-wide CRISPR loss-of-function and gain-of-function screens in T cells (*26–29*) onto the signalome (**Fig. 2B; Supplementary Table 2**). Across these datasets, 64 genes were identified in loss-of-function screens and 49 in gain-of-function screens. Genes implicated by CRISPR perturbation were significantly enriched for genes encoding SH2-domain-containing proteins (p = 2x10^-5^, Fisher’s exact test), underscoring their central role in T-cell signalling. Notably, 37 genes that were not transcriptionally regulated nonetheless exhibited functional effects in CRISPR screens, including seven core T-cell signalling genes encoding for SH2-domain-containing proteins (CBLB, GRB2, ZAP70, SLA2, CBL, LCP2, VAV1), whereas many transcriptionally regulated signalling genes were not detected in pooled CRISPR screens, highlighting potential gaps in existing perturbation-based maps (**Fig. 2A,B**).

**Figure 2:**
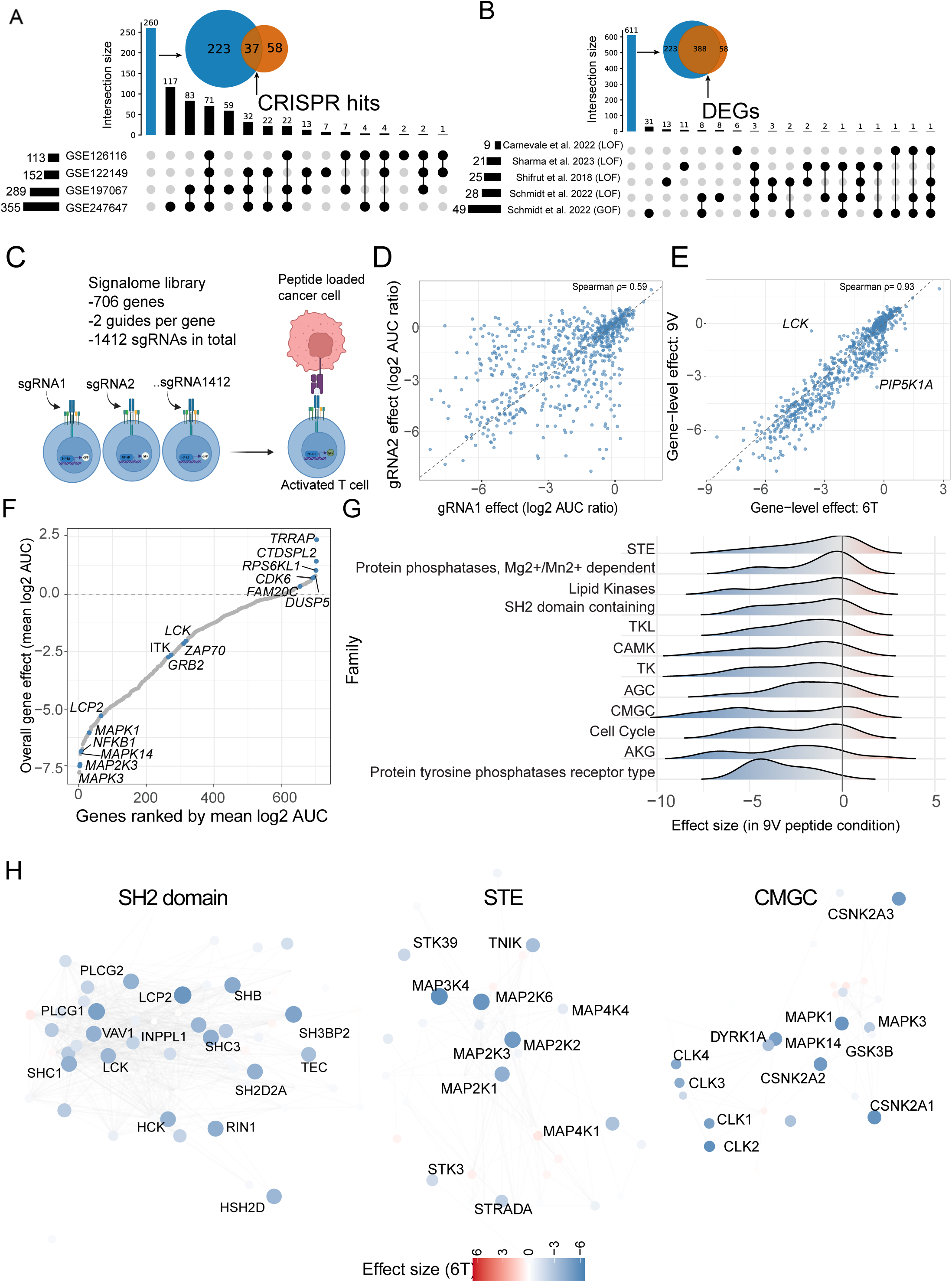
**A signalome-wide, arrayed CRISPR perturbation screen reveals hierarchical and distributed control of NF-κB signalling in T cells**. **A:** Overlap between genes that are not transcriptionally regulated across four T-cell–specific expression datasets and genes identified as hits in genome-wide CRISPR screens, revealing 37 signalling genes that are functionally required despite lacking dynamic expression changes. UpSet plot shows intersections across expression studies. **B:** Overlap between genes that are differentially expressed in at least one T-cell expression dataset and genes not identified as hits in genome-wide CRISPR screens, identifying 388 signalling genes that are transcriptionally regulated but not functionally required in pooled CRISPR screens. UpSet plot shows intersections across CRISPR studies. **C:** Schematic of the arrayed CRISPR perturbation strategy. **D:** Concordance between independent sgRNAs targeting the same gene under high-affinity stimulation (9V). Each point represents a gene; axes show log₂(AUC ratio) for sgRNA1 versus sgRNA2. **D:** Comparison of gene-level perturbation effects between low-affinity (6T) and high-affinity (9V) stimulation conditions. Each point represents one gene collapsed across the two sgRNAs. **F.** Genes ranked by overall effect size, defined as the mean gene-level log₂(AUC ratio) across both stimulation conditions. **G:** Distribution of effect sizes in 9V condition stratified by functional protein families **H:** Network representation of selected signalling modules with node colour indicating signed effect size under stimulation with low affinity antigen (6T). Nodes represent genes; edges correspond to curated functional interactions from STRING.

We next performed a signalome-wide, arrayed CRISPR perturbation screen to directly quantify the contribution of each component to NF-κB activation. 1G4-TCR⁺ Jurkat reporter cells were transduced with individual sgRNAs and stimulated through cell–cell co-culture with peptide-pulsed target cells. In total, ∼1,500 sgRNAs targeting ∼700 genes were assayed in 96-well format across 13 experimental batches (**Fig. 2C**). Each batch included internal controls, including an empty sgRNA vector and sgRNAs targeting known TCR signalling components (*LCP2, ITK*), enabling normalization of gene-level effects based on GFP signal area under the curve (AUC) relative to empty controls and an internal positive control to assess the editing efficiency (see details in **Methods**).

The screen showed high technical robustness. Independent sgRNAs targeting the same gene exhibited strong concordance, with ∼80% of guide pairs showing consistent effect direction (**Fig. 2D; Supplementary** **Figure. 2A**). Gene-level effects were highly correlated for T-cell stimulation with low-affinity (6T) and high-affinity (9V) antigens (Spearman ρ = 0.93), indicating that the core signalling architecture governing NF-κB activation is largely conserved across stimulation strengths (**Fig. 2E**). Most genes clustered near the identity line, whereas a subset showed context-dependent deviations. Notably, disruption of *LCK* strongly attenuated NF-κB activation under weak stimulation but had a markedly reduced effect under high-affinity stimulation (**Supplementary** **Figure. 2B**), consistent with partial bypass or compensation at higher signalling strength (*30*, *31*).

To characterise the global structure of signalling control, we ranked genes by their mean effect size across both conditions. The distribution was strongly skewed toward negative effects, consistent with depletion-driven signalling dependencies rather than enrichment effects (**Fig. 2F**). Canonical TCR signalling components, including *LCK, ZAP70, ITK, and LCP2*, ranked among the strongest negative regulators, validating the ability of this approach to recover core pathway architecture (**Fig. 2F**). TCR-specific signalling was among the highest-ranked pathways in the enrichment analysis, suggesting that the screen successfully recapitulated known genes while also assigning quantitative scores to signalome genes involved in regulating the NF-κB pathway (**Supplementary Figure 2C**).

We next examined how signalling control was distributed across protein families by comparing effect size distributions within each family (**Fig. 2G**). All families contributed to NF-κB regulation, but with distinct patterns. SH2-domain-containing proteins exhibited broad effect size distributions, indicating distributed engagement of multiple family members. In contrast, MAP kinase contributions were concentrated in a small number of nodes, leading towards ERK1/2 (*MAPK3/MAPK1*) and p38α (*MAPK14*) (**Fig 2F, H**). These patterns suggest that NF-κB signalling integrates both distributed adaptor-mediated control and more focal kinase bottlenecks, rather than being uniformly dominated by single components. Based on these quantitative dependencies, we reconstructed a data-driven network representation of NF-κB regulation, capturing hierarchical patterns of control, characterized by a small number of high-impact nodes alongside broadly distributed contributions from adaptor families (**Fig. 2H; Supplementary Table 3**).

Targeted perturbation of SH2-domain-containing proteins in primary CD8⁺ T cells validates functional TCR signalling dependencies Although large-scale perturbation screens with functional read outs are readily implemented in cell lines, extending such approaches to primary human T cells remains challenging. To enable arrayed perturbation in primary CD8⁺ T cells at scale, we established a workflow based on lentiviral delivery of sgRNAs followed by Cas9 electroporation, a strategy compatible with systematic perturbation, while avoiding the cost and scalability constraints of RNP-only approaches (*27*).

A further barrier to scalable functional assays in primary T cells is antigen specificity, as individual T cells respond only to their cognate TCR ligand. To circumvent this limitation, we employed bispecific molecules, which engage CD3 independently of endogenous TCR specificity (*32*, *33*). Here, the bispecific engager consists of an anti-CD3 single-chain variable fragment fused to a TCR targeting an HLA-A2-bound gp100 peptide (YLEPGPVTV), thereby triggering TCR signalling and directing T-cell activation against gp100-pulsed A375 melanoma cells (*18*). This approach enables the use of polyclonal primary CD8⁺ T cells while preserving physiologically relevant TCR-proximal signalling, thus substantially increasing experimental scalability.

Upon co-culture with gp100-pulsed target cells, polyclonal CD8⁺ T cells exhibited robust upregulation of activation markers (CD69, CD25, 4-1BB) and potent cytotoxicity (**Fig. 3A**). Despite these strong functional responses, phosphoprotein changes measured by CyTOF remained limited (**Fig. 3B, Supplementary Figure 3**), consistent with the low signalling flux observed during direct cell–cell interactions and reinforcing the sensitivity of functional readouts for capturing signalling outcomes (*18*).

**Figure 3:**
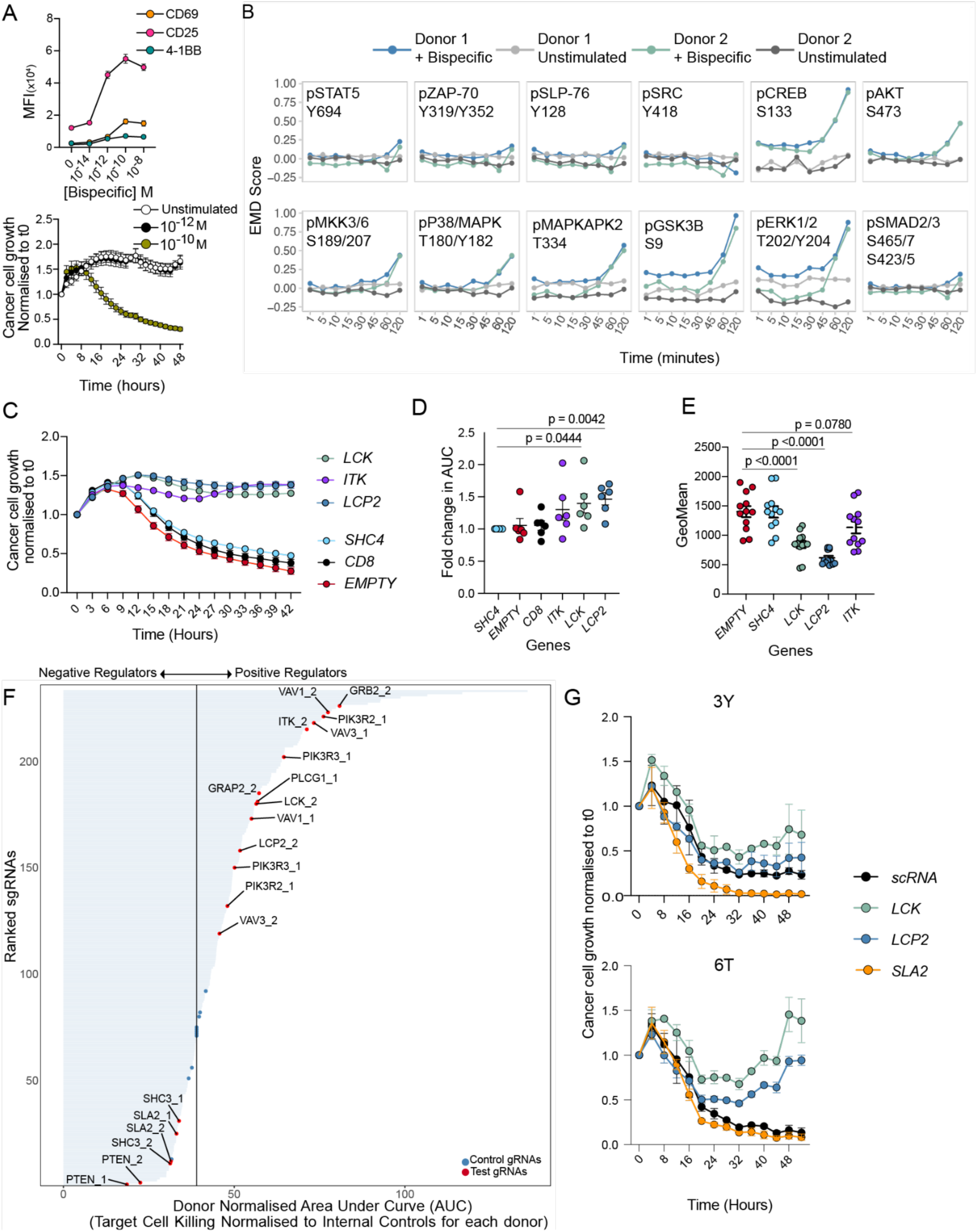
Targeted perturbation in primary CD8^+^ T-cells reveals the spectrum of activity of the SH2 gene family on T-cell cytotoxicity. A: Top panel - Expression (mean fluorescence intensity, MFI) of activation markers on CD8^-^ T-cells at 24 hours following co-culture with gp100 pulsed A375 melanoma cell lines and a bispecific T cell engager. Bottom panel - CD8^+^ cytotoxic responses following co-culture with A375 melanoma cell lines as described for the top panel. Data in both panels from 3 independent donors, representative of at least 3 experimental repeats. **B:** Phosphorylation of early signalling mediators downstream of TCR engagement as measured by mass cytometry (CyTOF). Data shown from two independent donors in one experiment, representative of at least 4 independent donors across 4 experiments. **C:** Representative killing curves from CD8^+^ T-cell co-cultures with A375 melanoma cells. T-cell activation was initiated through use of the bispecific engager at a final concentration of 10^-10^M. Targeting of known TCR-signalling mediators with sgRNAs inhibits cytotoxic T-cell responses. Data show from one donor and one independent experiment, representative of at least 6 independent donors across 6 independent experiments. **D:** Area under the curve (AUC) analysis of killing curves shown in panel C, normalised to SHC4 targeted controls. Data from 6 independent donors across 6 independent experiments. Statistical analysis performed by paired T-test. **E:** Targeting TCR-signalling mediators with sgRNAs reduces surface 4-1BB expression, measured at 48 hours post co-culture with A375 melanoma cells as detailed in D. Data shown are the same T-cells from panel D, and are from 6 independent CD8^+^ donors across 6 independent experiments. Statistical analysis performed via One-Way ANOVA with Dunnett’s multiple comparison tests. **F:** Waterfall plot of normalised AUCs from killing curves illustrated in panel C, from experiments targeting all genes encoding SH2-domain containing proteins. Killing curves for each perturbation were first normalised to donor-specific internal negative control sgRNAs before computing AUC values. Ranked gene analysis identifies both positive and negative regulators of T-cell cytotoxicity within the SH2 gene family. **G:** Representative killing curves from targeting *LCK*, *LCP2* or *SLA2* in primary 1G4-TCR^+^ CD8^+^ T-cells. A375 melanoma cell lines were pulsed with NY-ESO-1 peptides as indicated to final concentrations of 100µg/mL. Data shown from one independent experiment, representative of two independent repeats using two individual CD8^+^ donors.

To validate the signalling architecture inferred from the Jurkat signalome screen in a physiological setting, we focused on the SH2-domain-containing protein family, which exhibited broad functional engagement in NF-κB regulation despite lacking a coherent pattern of transcriptional regulation during T-cell activation (**Fig. 2G**). This family includes multiple well-established TCR signalling components, making it a suitable test case for functional validation. We targeted 66 SH2-domain-containing genes, along with control targets, in polyclonal primary CD8⁺ T cells from six independent donors using lentiviral sgRNA delivery followed by Cas9 electroporation (**Supplementary Figure 4A**). Functional consequences were assessed by quantifying target cell killing in co-culture assays (**Fig. 3C**).

Disruption of canonical TCR signalling components, including *LCK*, *LCP2*, and *ITK*, markedly impaired CD8⁺ T-cell cytotoxicity (**Fig. 3C,D**) and reduced expression of activation markers 4-1BB (**Fig. 3E**) and CD25 (**Supplementary Figure 4B**) at 48 hours, validating the sensitivity and specificity of the assay. After correcting for donor and batch effects (**Methods**), systematic perturbation of the SH2 gene family revealed a spectrum of functional effects on cytotoxicity **(Fig. 3F)**.

Genes were ranked by their impact on tumour cell killing, with cytotoxicity levels quantified using the AUC metric (**Methods**). Our analysis revealed profiles that closely mirrored both known TCR signalling annotations, and the dependencies observed in the Jurkat screen (**Fig. 3F**). Perturbation of key signalling mediators (*ITK*, *LCK*, *LCP2*, *PLCγ*, *PIK3R1-3*), adaptor proteins (*GRAP*, *GRB2*), and cytoskeletal regulators (*VAV1*, *VAV3*) consistently reduced CD8⁺ T-cell cytotoxicity, whereas disruption of *SLA2* and *PTEN*, both known negative regulator of TCR signalling, enhanced CD8⁺ T-cell-mediated killing (**Fig. 3G; Supplementary Fig. 4C,D**). These results demonstrate that the SH2 gene family constitutes a distributed and functionally essential regulatory module in primary T-cell activation.

Because bispecific-based activation bypasses endogenous antigen specificity, we next asked whether these dependencies were preserved in antigen-specific TCR-engineered T cells. Primary CD8⁺ T cells were transduced with plasmids encoding the 1G4-TCR and an LNGFR tag, and TCR Vβ13.1⁺ LNGFR⁺ cells were sorted and expanded for use *in vitro* (**Supplementary Fig. 4E**). This system enabled direct comparison of signalling requirements across different ligand affinities. Consistent with the Jurkat screen, disruption of *LCK* exhibited affinity-dependent effects: under high-affinity stimulation with the 3Y peptide, *LCK* targeting had minimal impact on tumour cell killing, whereas under weaker stimulation conditions (6T), cytotoxicity was strongly impaired (**Fig. 3G**).

Together, these results demonstrate that functional signalling dependencies identified in Jurkat cells are largely conserved in primary human CD8⁺ T cells. This validation establishes the robustness of the perturbation-based framework across experimental systems and highlights how distributed adaptor-mediated regulation and signal-strength–dependent buffering shape TCR-driven functional responses.

## *TRRAP* and *CTDSPL2* constrain CD8⁺ T-cell effector output downstream of TCR signalling

Although the majority of perturbations in the signalome screen reduced NF-κB activity, a small number of gene disruptions increased reporter output. Among these, targeting of *TRRAP* and *CTDSPL2* consistently enhanced NF-κB-driven GFP expression in Jurkat cells (**Fig. 4A**). TRRAP is a pseudokinase that functions as a scaffold within histone acetyltransferase complexes involved in transcriptional regulation, whereas CTDSPL2 is a poorly characterized serine/threonine phosphatase. Neither protein has been well defined in the context of T-cell signalling, prompting further functional validation in primary cells.

**Figure 4:**
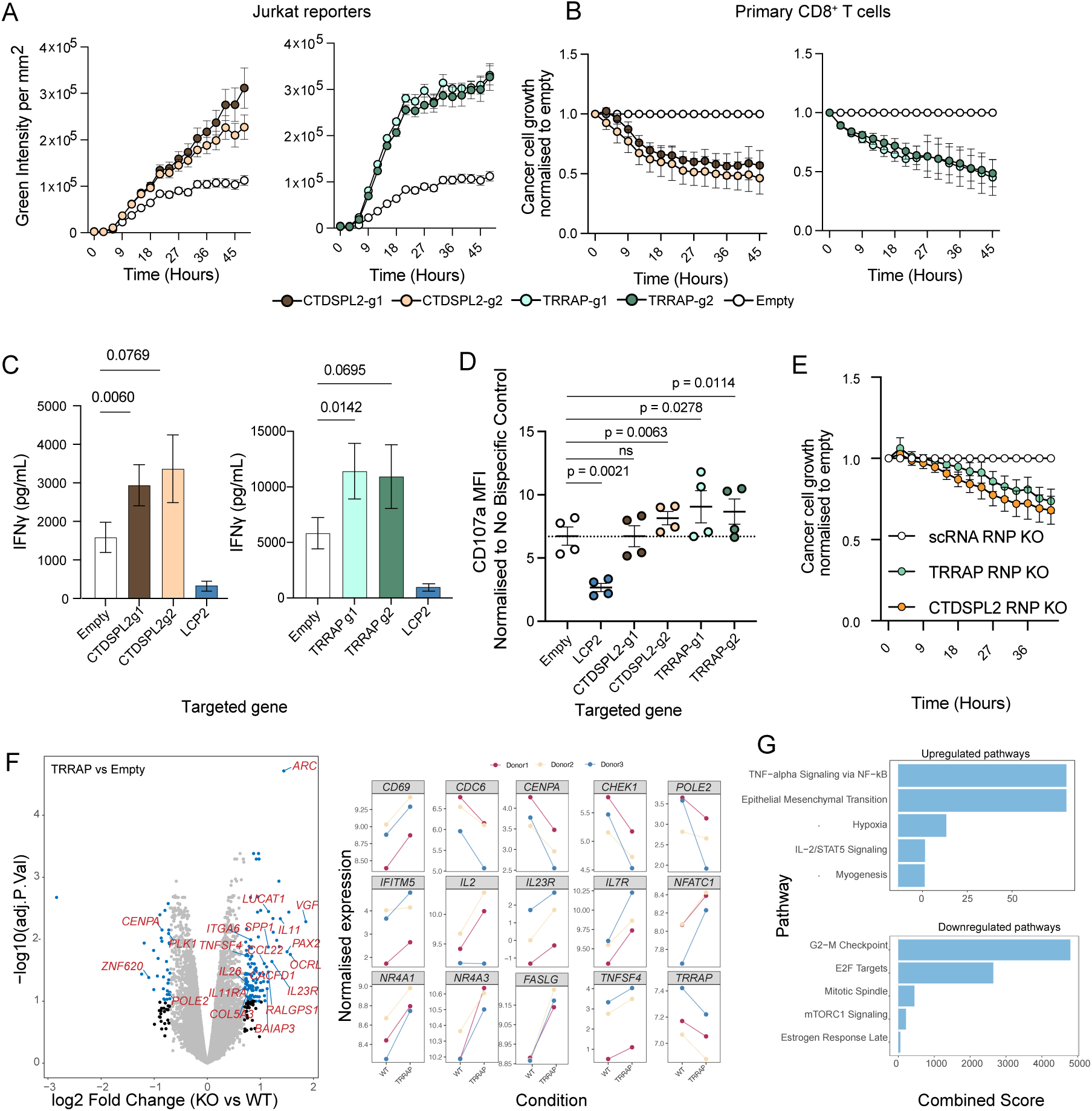
Perturbation of novel targets *CTDSPL2* and *TRRAP* enhances primary CD8^+^ T-cell functional activity. A: NFkB-GFP traces from 1G4-TCR^+^ Jurkat reporters when transduced with *CTDSPL2* (left) or *TRRAP* targeting sgRNAs (right). Jurkat cells were stimulated with A375 melanoma cell lines pulsed with NY-ESO-1 9V peptide (100µg/mL). **B:** Cytotoxic responses from primary CD8^+^ T-cells when co-cultured with A375 melanoma cell lines pulsed with gp100 peptide and a bispecific T-cell engager (final concentration 10^-11^M). Data shown from 9 independent donors across 3 experiments (TRRAP sgRNAs) or a minimum of 4 independent donors across 2 experiments (CTDSPL2 sgRNAs). A375 cell growth was normalised to the empty control for each donor. **C:** IFNᵧ secretion at 24 hours from primary CD8^+^ T-cells stimulated as described for panel B. Data shown from 2 independent donors across 1 individual experiment, representative of at least 4 independent donors across 2 experiments. Statistical analysis performed via One-Way ANOVA with Dunnett’s multiple comparisons. **D:** CD107α mean fluorescence intensity (MFI) from CD8^+^ T-cells stimulated as described for panel B, measured at 4 hours post stimulation. CD107a expression was normalised to unstimulated controls (no bispecific) for each CRISPR targeting condition. Data shown from 4 independent donors across 2 experiments. **E:** Cytotoxic responses of 1G4-TCR^+^ primary CD8^+^ T-cells when co-cultured with A375 melanoma cell lines expressing basal NY-ESO-1. 1G4-TCR^+^ CD8^+^ T-cells were electroporated with RNPs targeting either *CTDSPL2* or *TRRAP*, together with a scrambled control (scRNA). Data shown from 6 donors across 3 independent experiments. Growth curves were normalised to the scRNA controls for all donors. **F.** Differential gene expression analysis of *TRRAP*-targeted primary CD8⁺ T cells following stimulation. Volcano plot showing log₂ fold change (KO vs WT) versus –log₁₀ adjusted P value from donor-controlled limma-voom analysis (n = 3 donors). Selected representative genes are highlighted. Inset panels show donor-resolved normalized expression for representative upregulated inflammatory genes (e.g., IL2, IL7R, IL23R, TNFSF4) and downregulated proliferative genes (e.g., CDC6, CENPA, CHEK1, POLE2). Lines connect paired donor samples. TRRAP transcript abundance was modestly but consistently reduced across donors (see Fig. 4F, inset). **G**. Pathway enrichment analysis of differentially expressed genes in *TRRAP*-targeted cells. Hallmark gene set enrichment (Enrichr) identified suppression of proliferative programs, including G2/M checkpoint and E2F target pathways, and enrichment of inflammatory and TNFα–NF-κB signalling modules. Combined enrichment scores are shown.

To assess their roles in physiological settings, we targeted *TRRAP* or *CTDSPL2* in primary human CD8⁺ T cells using lentiviral sgRNA delivery followed by Cas9 electroporation.

Disruption of either gene significantly enhanced CD8⁺ T-cell-mediated killing of A375 melanoma cells in bispecific-based co-culture assays (**Fig. 4B**). Consistent with increased effector activity, targeting *TRRAP* or *CTDSPL2* also elevated IFNγ secretion at 24 hours (**Fig. 4C**). In addition, after targeting *TRRAP* or *CTDSPL2* we observed enhanced degranulation, as measured by surface CD107α expression at 4 hours post A375 co-culture (**Fig. 4D**). These results indicate that both proteins act to constrain T-cell effector responses downstream of TCR-proximal signalling.

Because bispecific-based activation bypasses endogenous antigen specificity, we next examined whether these effects were preserved in antigen-specific TCR-engineered T cells. Primary CD8⁺ T cells expressing the 1G4 TCR were electroporated with sgRNA–Cas9 ribonucleoprotein (RNP) complexes targeting *TRRAP* or *CTDSPL2*, and cytotoxic activity was assessed against A375 melanoma cells presenting endogenous levels of NY-ESO-1. Importantly, these experiments utilised a distinct, non-overlapping sgRNA sequence compared to these utilised in our signalome-wide arrayed screen. Under these basal antigen conditions, disruption of either gene significantly enhanced tumour cell killing relative to scrambled controls (**Fig. 4E**). This enhancement was maintained across a range of peptide affinities, again utilising 6T or 3Y NY-ESO-1 altered peptide ligands (**Supplementary Fig. 5A**).

Finally, to assess whether enhanced CD8^+^ T-cell effector functions arising from *TRRAP* or *CTDSPL2* perturbations were accompanied by an increase in markers of T-cell dysfunction and exhaustion (e.g., PD-1, LAG-3 and TIM-3), cell surface expression of these molecules was assessed in polyclonal CD8^+^ T-cells 24 hours post stimulation with A375 melanoma cells and the bispecific engager. Phenotypically, *TRRAP* or *CTSPL2* targeted T-cells were similar to the unedited, empty plasmid controls, where expression of these three inhibitory receptors displayed an activation-marker-like expression pattern. Expression of surface PD-1, LAG-3 and TIM-3 all correlated with the bispecific concentration utilised to initiate T-cell activation in these experiments, and reduced expression was observed in the *LCP2* targeted control (**Supplementary Fig. 5B**).

To investigate how TRRAP and CTDSPL2 depletion enhanced cytotoxicity, we performed RNA sequencing following CRISPR-mediated perturbation in primary CD8^+^ T-cells. TRRAP is a scaffold component of the SAGA and TIP60 histone acetyltransferase complexes and supports MYC- and E2F-dependent transcriptional programs. Consistent with this role, upon activation, TRRAP-deficient T cells exhibited coordinated downregulation of genes associated with cell cycle progression and DNA replication, including CENPA, PLK1, POLE2, and multiple MCM complex components. In contrast, inflammatory and immune signalling genes, including IFITM5, IL23R and NR4A family components were upregulated (**Figure 4F, Supplementary Table 4**). Pathway-level analysis revealed suppression of proliferative programs alongside enrichment of inflammatory and cytokine signalling modules (**Figure 4G**). These findings suggest that TRRAP loss shifts T cells away from a proliferative transcriptional state toward an inflammatory, effector-associated profile. By contrast, CTDSPL2-deficient cells displayed minimal global transcriptional remodelling. Donor-paired analyses demonstrated that most genes remained near baseline at both stimulated and unstimulated conditions (**Supplementary Fig. 5C, Supplementary Table 4**). The increased killing arising from *CTDSPL2* perturbation is, therefore, unlikely to arise from large-scale transcriptional reprogramming and instead reflects altered signalling dynamics.

Together, these data identify TRRAP and CTDSPL2 as previously unrecognized negative regulators of CD8⁺ T cell effector function. Their disruption enhances cytotoxicity, cytokine production, and degranulation under both polyclonal and antigen-specific activation conditions without inducing excessive expression of inhibitory receptors. These findings highlight novel regulatory nodes that may be therapeutically leveraged to enhance T cell–based immunotherapies.

## Discussion

In this study, we present a perturbation-based framework to interrogate cellular signalling during physiological cell–cell engagement. Leveraging functional readouts rather than direct measurement of signalling intermediates allowed us to identify key signalling regulators despite remarkably low signalling flux under these conditions (*18*). By quantifying transcription factor activity following systematic perturbation of signalling components, we could capture pathway activity through integrated transcriptional outputs rather than discrete molecular states. This approach is motivated by models of temporal signal integration, as articulated by Purvis and Lahav, and by subsequent optogenetic studies demonstrating that many proximal T-cell signalling intermediates decay rapidly following receptor disengagement (*34*, *35*). In contrast, transcriptional outputs integrate signalling over time, accumulating to provide a longer-lived functional record of signalling activity. In the context of T-cell activation, our findings support this framework by demonstrating that robust NF-κB activation and effector responses results from low-amplitude or spatially restricted signalling during direct T-cell-tumour cell engagement.

A key feature of this framework is its compatibility with physiologically relevant stimulation formats. By interrogating signalling in cell–cell co-culture systems, our approach avoids reliance on supraphysiological stimulation commonly required to elicit detectable changes in phosphorylation or kinase activity, and instead captures the integration of complex signal inputs that arise during direct cellular interactions. Although strong stimulation (most often achieved using anti-CD3/CD28 bead-based activation) has been instrumental for defining canonical signalling pathways in global phosphoproteomic studies (*3*, *5*, *11*), it may not reflect signalling behaviour in physiological settings. This limitation is particularly relevant in studies of T-cell activation, where signalling outcomes reflect the combined influence of antigen affinity, co-stimulatory, and inhibitory inputs, such that maximal pathway activation cannot be assumed to be an appropriate starting point for interrogating authentic signalling processes. Similar considerations apply to high-dimensional single-cell approaches, including mass cytometry, which have provided powerful insights when phosphorylation flux is sufficiently robust, but which typically rely on maximal signalling to infer signalling relationships (*17*, *36*).

Applying this strategy to T-cell receptor signalling revealed that the nodes important for functional signalling are largely unchanged at different stimulation strengths produced by differing-quality TCR ligands, whereas there do exist specific nodes that are highly context dependent. Perturbation of *LCK*, a widely studied SRC-kinase which is known to initiate receptor signalling from the TCR (31) exhibited a strong dependence on ligand affinity; disruption of both markedly impaired NF-κB activation under weak stimulation but was buffered under high-intensity conditions. Quantitative modulation of proximal kinase activity has been described for LCK. Partial reduction of LCK activity preferentially impairs T cell responses to low-affinity or low-dose antigen, whereas strong stimulation can partially compensate for diminished kinase availability (*30*, *37*, *38*). Notably, Cabezas-Caballero and colleagues demonstrated that altering coreceptor identity (swapping CD8 and CD4) effectively redistributes and sequesters LCK, thereby tuning TCR sensitivity by modulating the local availability of active kinase. This work highlights how LCK availability functions as a quantitative regulator of signalling output. Together, these findings outline how targeting key proximal signalling components can serve as a mechanism to engineer tuning of cellular signalling in the context of varied stimulation strengths.

Beyond individual nodes, our data reveal how signalling control is unequally distributed across molecular families. Rather than treating pathway components as interchangeable, our perturbation-based analysis quantifies family-level engagement, distinguishing signalling nodes dominated by a small number of high-impact components from those that rely on distributed contributions. This is exemplified by the MAPK pathway, in which NF-κB regulation was driven primarily by a limited set of kinases, consistent with bottlenecked enzymatic cascades. In contrast, SH2-domain–containing proteins exhibited broad functional engagement, reflecting their role as proximal adaptors that integrate receptor-derived signals and connect them to diverse downstream pathways. These patterns are consistent with the early proposal by Pawson and colleagues, i.e., that signalling networks are assembled from conserved interaction modules whose combinatorial deployment confers specificity, robustness, and evolutionary adaptability without reliance on rigid, linear pathways (*39–41*). Our inferred signalling network therefore supports a modular, hierarchical organisation in which control is unevenly distributed across molecular layers and funnelled through preferred pathways under defined conditions.

Notably, many SH2-domain-containing proteins exerted strong functional effects despite limited transcriptional regulation across T-cell activation states. This suggests that signalling control at this level operates largely through constitutively expressed components whose regulatory capacity is governed post-translationally, through phosphorylation, conformational changes, and dynamic recruitment to signalling complexes. As a result, expression-based analyses may underestimate this contribution to signalling control, whereas perturbation-based functional approaches are particularly well suited to reveal their role.

Finally, our experimental framework enabled the identification of previously unrecognised negative regulators of T cell effector function, specifically TRRAP and CTDSPL2. TRRAP functions as a core scaffold for multiple histone acetyltransferase complexes, thereby regulating chromatin accessibility and transcriptional output downstream of diverse signalling pathways (*42*). Consistent with this role, TRRAP depletion resulted in coordinated suppression of proliferative and DNA replication–associated transcriptional programs alongside enrichment of inflammatory and immune signalling modules, indicating a redistribution of transcriptional state rather than global activation. In contrast, CTDSPL2 is a poorly characterised serine/threonine phosphatase implicated in transcriptional regulation and cell-state control, and its role in T cell biology has not previously been examined (*43*). Notably, CTDSPL2 depletion enhanced cytotoxicity with minimal global transcriptional remodelling, suggesting that its effects are mediated primarily through altered signalling dynamics rather than broad transcriptional reprogramming. These findings provide functional validation of nodes identified in the signalome-wide arrayed perturbation screen and illustrate how this approach can uncover regulatory layers that modulate signalling output and effector potential that may be overlooked by pooled genetic screens.

In conclusion, whereas genome-wide perturbation screens and high intensity stimulation-based approaches have been effective for defining essential signalling components, they are less well-suited to dissecting signalling networks dependent on quantitative effects. By emphasising quantitative, time-resolved functional outputs in the setting of physiological stimulation, the framework presented here offers a general strategy for interrogating cell responses as the field moves toward a system-level understanding of signalling.

## Materials and Methods

### Cell culture

All Jurkat cell lines were cultured in RPMI-1640 medium (Gibco, #21875-034) supplemented with 10% FCS (Gibco, #A5256801, Lot #B3024242RP), 1% Penicillin-Streptomycin-Neomycin (Sigma-Aldrich, #P4083), 1% L-Glutamine (Sigma-Aldrich, #G7513), 1% Sodium Pyruvate (Sigma-Aldrich, #S8636) and 1% HEPES (Gibco, #15630-056). HEK293-T cells for lentivirus production were cultured in DMEM complete medium (Gibco, #41965-039) supplemented as described for the RPMI-1640 Jurkat medium. A375 melanoma cell lines were cultured in DMEM-F12 medium (Gibco, #11320-033) supplemented with 10% FCS, 1% Penicillin-Streptomycin-Neomycin, 1% Sodium Pyruvate and 1% HEPES. Finally, all primary human cells were cultured in RPMI-1640 medium supplemented with 10% human serum, 1% Penicillin-Streptomycin-Neomycin, 1% L-Glutamine, 1% Sodium Pyruvate, 1% non-essential amino acids (NEAA, Gibco, #11140), 1% HEPES and 0.1% Β-mercaptoethanol (Gibco, #31350-010).

### Production of 1G4^+^ Jurkat NF-kB GFP reporter cell lines

Jurkat T-cells (ATCC) were first transduced with the Cas9 encoding plasmid pKLV2-EF1a-Cas9Bsd-W, as previously described (*26*), to produce Cas9 expressing Jurkat cell lines. Next, Cas9^+^ Jurkat cells were transduced with pSIRV-NF-kB-eGFP construct which was a gift from Peter Steinberger (Addgene plasmid # 118093)(*44*) prior to single cell cloning and selection of NF-kB GFP expressing clones. Endogenous TCR was then deleted from these lines via electroporation with a TRAC/TRBC sgRNA - Cas9 RNP complex. For the generation of RNPs, sgRNAs targeting TRAC and TRBC1/2 (IDT, TRAC sequence: AGAGTCTCTCAGCTGGTACA, TRBC1/2 sequence: GGAGAATGACGAGTGGACCC) (*45*) was pre-incubated with Cas9 (IDT, #10007807) at room temperature for 30 minutes. Here, 0.73µL of each sgRNA (400µM) was added to 0.5µL of Cas9 (62µM). Electroporation was then performed using the Lonza P3 Nucleofector kit (Lonza, # V4SP-3096) as per manufacturer’s instructions. Briefly, one million Jurkat cells were resuspended in 20µL P3 buffer (Lonza, #V4SP-3096), to which the complexed RNP was added. Cells were then moved to one well of a 16-well electroporation strip for electroporation with a Lonza Nucleofector on programme #E0115. Cells were immediately recovered from the electroporation strip by the addition of 80µL of media before returning them to culture. Following deletion of endogenous TCR, Cas9^+^ NF-kB GFP^+^ TCR^-/-^ Jurkats were then transduced with constructs encoding for the 1G4-TCR (*18*) to produce the final 1G4^+^ Cas9^+^ NF-kB GFP^+^ Jurkat cell lines.

### sgRNA design and cloning

To design sgRNA molecules, the top two candidates from CRISPick (Broad Institute) were selected for each gene. sgRNA sequences were purchased as oligos in both forward and reverse orientations (Sigma-Aldrich) prior to annealing, and were then cloned into the BbsI site of the CRISPR sgRNA expression vector pKLV2-U6gRNA5(BbsI)-PGKpuro2ABFPW (Addgene, #67974) as previously described (*46*). All sgRNA sequences produced for the signalome library are displayed in supplementary table 1.

### Lentivirus production

Lentiviruses were prepared by transfecting HEK293T with the sgRNA expression vector. Firstly, 500ng sgRNA expression plasmid was combined with 250ng pMD2.G (Addgene, #12259) and 1.2µg psPAX.2 (Addgene, #12260) in 500µL optiMEM (Gibco, #13985062). To this, 2µL plus reagent (Thermo Fisher, #15338100) was added prior to mixing by inversion and incubation at room temperature for 5 minutes. Next, 6µL lipofectamine-LTX (Thermo Fisher #15338100) was added and the solution was again inverted to mix, prior to incubation at room temperature for 30 minutes. This optiMEM mixture was then added to HEK293T cells in a 6 well plate together with 2mL of DMEM medium supplemented as described above. Lentiviral supernatants were then harvested at 48-72hrs and filtered through a 0.45µm filter.

All lentiviruses were either used immediately or stored at -80°C until use, but were never freeze-thawed.

### Peptide sequences for A375 pulsing

Where indicated, A375 melanoma cell lines were pulsed with the following NY-ESO-1 altered peptide ligands to induce TCR signalling downstream of the 1G4-TCR. Peptide pulsed A375 cell lines were co-cultured with either Jurkat or primary CD8^+^ T-cells as indicated, and final peptide concentrations are shown throughout the text. All peptides were ordered from Genscript with a purity >95% and were reconstituted in DMSO upon arrival, stored at -80°C, and never freeze-thawed more than twice. 9V: SLLMWITQV, 3Y: SLYMWITQV, 3I: SLIMWITQV, 6T: SLLMWTTQV. Affinities (K_D_ for these NY-ESO-1 peptides have been previously described (9V ≌ 72μM, 3Y ≌ 30μM, 3I ≌ 17μM, 6T ≌ 101μM (*47*). Finally, for experiments utilising bispecific T-cell engagers, A375 cell lines were pulsed with the gp100 peptide (10µg/mL, sequence: YLEPGPVTV) as described, which was also obtained from Genscript at >95% and handled as per the NY-ESO-1 altered peptide ligands.

### Arrayed screening of the signalome in 1G4-TCR^+^ Jurkat reporter cell lines

1x10^5^ 1G4-TCR^+^ Jurkat NF-kB GFP reporter cells were transduced with 100µL of the arrayed library lentivirus in a 96-U bottomed plate, in a total of 200µL. Transduction was performed by centrifugation at 800g, 90min, 32°C. After 48hrs, transduced cells were grown in the presence of puromycin (1µg/mL final concentration, Gibco, #A11138-03) to select only transduced cells. At day 7, transduced Jurkat reporter cells were counted via flow cytometry alongside simultaneously ensuring transduction and selection success via expression of the BFP tag in the sgRNA plasmid. All cells were 95-100% BFP^+^. Following this counting, all culture volumes across the arrayed library were normalised to allow for consistent cell numbers to be utilised in co-cultures with A375 melanoma cells. Here, A375 melanoma cells were treated with 100µg/mL of 6T or 9V NY-ESO-1 peptides (alongside a no peptide control), before plating 5,000 cells in 25µL in a 384 well plate (Falcon, #353961), to which 10,000 Jurkat reporter cells were added in 25µL. NF-kB signalling pathway activation downstream of TCR engagement was measured using IncuCyte imaging for 42-48 hours.

### Arrayed screen analysis in Jurkat cells

Raw IncuCyte time-course data from multiple experiments and plates were aggregated and annotated with experimental metadata. Condition labels were harmonized, and control wells were explicitly annotated. For each peptide, signalling output was summarized by calculating the area under the curve (AUC) using trapezoidal integration. Data was collected for a total of 706 genes with signalome library. AUC values were normalized within each experiment and condition to the median AUC of empty control wells and log2-transformed for downstream analysis.

Each gene was targeted by two independent gRNAs. Guide-level effects were computed as mean log2-normalized AUC ratios across replicates. Gene-level effects were derived using a rule-based guide collapsing strategy that accounts for guide concordance and effect strength. When guide effects were concordant (AUC within 1), their median was used; in cases of strong discordance (particularly when one guide showed a weak effect or when guides had opposite signs) the guide with the larger absolute effect was retained. Guide concordance was assessed using Spearman correlation and same-direction agreement.

Gene-level effects were computed independently for each stimulation condition (6T and 9V) and compared using Spearman correlation. A composite discordance score integrating effect size differences and sign reversals was used to prioritize condition-specific dependencies. Genes were annotated with functional families and essentiality, and family-level effect distributions were visualized to identify pathway-level trends.

### Primary CD8^+^ T-cell transduction

Primary CD8^+^ T-cells were isolated from healthy donor leukocyte cones obtained from the NHS blood-transfusion service. CD8^+^ T-cells were isolated using RosetteSep human CD8^+^ T-cell enrichment cocktail (StemCell, #15023) as per manufacturer’s instructions. Isolated CD8^+^ T-cells were then stimulated for 48 hours prior to transduction with 10µL/mL of CD3/CD28 immunocult (StemCell, #10971) and 50U/mL IL-2 (Biolegend, #589104), with cells at 2x10^6^/mL and 2mL/well in a 48 well plate. 250µL of Retronectin solution (Takara Bio, #351178) at 25µg/mL was added to non-treated 24-well tissue culture plates (Costar, #3738) for 24 hours at 4°C prior to transduction. On the day of transduction, Retronectin solution was removed and plates were blocked by addition of 300µL of 2% BSA in PBS (w/v) for 30 minutes at room temperature. The BSA solution was then removed and wells were washed once in 500µL PBS, before addition of 1.5mL of lentiviral supernatant and centrifugation at 2000g for 90 minutes, 32°C, no brake. Following centrifugation, all lentiviral supernatant (except 200-250µL to prevent the well from drying) was removed from the well and 10^6^ CD8^+^ T-cells were seeded into the well in 1mL of complete RPMI with supplementation of IL-2 at 50U/mL. Cells were briefly spun onto the lentiviral coated wells at 2000rpm for 2 minutes, 32°C with the brake applied. Transduction efficiency was assessed 48-72 hours post transduction by measuring BFP expression via flow cytometry. Transduced cells were then selected by the addition of puromycin to 1µg/mL (Gibco, #A11138-03), prior to electroporation with Cas9 protein.

### Electroporation of primary CD8^+^ T-cells

Transduced primary CD8^+^ T-cells were electroporated with Cas9 protein (IDT, #10007807) once BFP expression had reached >75-80%. Electroporation was performed using the Lonza P3 Primary cell nucleofection kit (Lonza, #V4SP-3096) as per manufacturer’s instructions. Here, 10^6^ CD8^+^ T-cells were resuspended in 20µL P3 buffer (Lonza, #V4SP-3096), to which 0.5µL Cas9 (62µM stock) was added. CD8^+^ T-cells were then transferred to 1 well of the 16-well electroporation strip before electroporation on programme #E0115. Immediately after electroporation, cells were recovered by the addition of 80µL of complete RPMI-1640 medium supplemented with 50U/mL IL-2. Cells were then plated across 2 wells in a 96 well-flat bottomed tissue culture plate, each in 200µL with IL-2 at 50U/mL. Cells were maintained at 1-2x10^6^/mL, in the presence of 50U/mL IL-2 and 1µg/mL puromycin, until day 7 post electroporation at which point functional assays were performed. All sgRNA sequences used in the signalome library are displayed in supplementary table 1. Where indicated, perturbations were also performed with RNPs, using independent and non-overlapping guides (IDT) from the signalome library. Guide sequences used for RNP based perturbations are as follows: TRRAP: AGTGGTCTGGTCAACCACCG, CTDSPL2: TTATTCATCAGCCCACGCGG, LCK: GACCCACTGGTTACCTACGA, LCP2: TGAAGAAGTACCACATCGAT, SLA2: GGGACCGGATCAGACACTAC, scRNA: GTATTACTGATATTGGTGGG.

### A375 melanoma cytotoxicity and T-cell degranulation assays

At day 7 post Cas9 electroporation, 5x10^4^ A375 mOrange^+^ melanoma cells were plated in 100µL in a 96 well-flat bottomed tissue culture plate and pulsed with gp100 peptide at a final concentration of 10µg/mL. Cells were incubated at 37°C for at least 4 hours to allow for cell adherence and antigen presentation. A375 cells were then washed once in PBS before addition of 1x10^5^ CD8^+^ T-cells in 100µL, and 100µL of the gp100-HLA-A2 - CD3 bispecific engager to final concentrations as indicated. Details of this bispecific T-cell engager have been previously described (*18*). Growth of the A375 melanoma cell line was then tracked for approximately 48 hours via IncuCyte imaging, tracking the A375-expressed mOrange fluorescent protein, allowing for quantification of T-cell mediated cytotoxicity (*26*). CD8^+^ T-cell degranulation was assessed at 4 hours post co-culture by flow cytometry with staining with the following antibodies: CD8 PeCy7, 1:200, Biolegend #344712, CD107α PE, 1:500, Biolegend #328608 and eBioscience fixable viability dye 780, 1:1000, Thermo Fisher #65-0865-18.

### RNA sequencing in primary CD8^+^ T-cells

To assess transcriptional changes with *TRRAP* perturbation, we first targeted *TRRAP* via lentiviral delivery of sgRNA and electroporation of Cas9 protein as previously described. For these experiments, three primary CD8^+^ T-cell donors were transduced with lentiviruses encoding the TRRAPg1 sequence, or empty plasmid controls. On day 7 post Cas9 electroporation, 5x10^5^ CD8^+^ T-cells were activated on OKT3 coated 48-well tissue culture plates for 3 hours. OKT3-coated tissue culture plates were prepared in advance, where each well was treated with 300µL of Ultra-LEAF OKT3 at 1µg/mL (Biolegend, #317326), or PBS only for unstimulated controls. After 3 hours, cells were recovered, washed in PBS and RNA was extracted using the Qiagen RNA mini kit (#74104) as per manufacturer’s instructions.

Isolated RNA was then sent to Novogene for NGS. FASTQ files were aligned using STAR version 2.7.3a and quantified using featureCounts version 2.0.6. Ensembl Homo sapiens GRCh38.86 was used as the reference genome. Lowly expressed genes were filtered out, and gene expression distributions of each sample were normalised using edgeR version 3.42.2. Differential expression analysis was performed using limma version 3.54.2.

For experiments investigating transcriptional changes associated with *CTDSPL2* perturbation, primary CD8^+^ T-cells from two donors were transduced with lentiviruses encoding for the CTDPSL2g1 or CTDSPL2g2 sequences, alongside empty plasmid controls. Again, at day 7 post Cas9 electroporation, T-cells were stimulated as described as per *TRRAP* perturbations. At 3 hours post stimulation on plate-bound OKT3, T-cells were collected and resuspended in 50µL DNA/RNA shield (Zymo Research, #R1100-250) and sent to Plasmidsaurus for NGS. Data handling and analysis was performed by Plasmidsaurus as follows: Quality of the fastq files was assessed using FastQC v0.12.1.

Reads were then quality filtered using fastp v0.24.0 with poly-X-tail trimming, 3’ quality-based tail trimming, a minimum Phred quality score of 15, and a minimum length requirement of 50bp. Quality-filtered reads were aligned to the reference genome using STAR aligner v2.7.11 with non-canonical splice junction removal and output of unmapped reads, followed by coordinate sorting using samtools v1.22.1. PCR and optical duplicates were removed using UMI-based deduplication with UMIcollapse v1.1.0. Alignment quality metrics, strand specificity, and read distribution across genomic features were assessed using RSeQC v5.0.4 and Qualimap v2.3, with results aggregated into a comprehensive quality control report using MultiQC v1.32. Gene-level expression quantification was performed using featureCounts (subread package v2.1.1) with strand specific-counting, multi-mapping read fractional assignment, exons and three prime UTR as the feature identifiers, and grouped by gene_id. Final gene counts were annotated with gene biotype and other metadata extracted from the reference GTF file.

### Flow cytometry for assessment of CD8^+^ T-cell phenotype

Genetically perturbed CD8^+^ T-cells were stimulated on gp100 pulsed A375 melanoma cell lines together with the bispecific T-cell engager as described. At 24 hours post stimulation, T-cells were recovered and transferred to a 96-U bottomed plate before staining with 100µL of antibody mix containing the following antibodies, all used at 1:200 unless otherwise stated: CD8 PE-Cy7, Biolegend, #344750, PD-1 Alexa Fluor 647, Biolegend, #329910, LAG3 FITC, Biolegend, #369308, TIM-3 PE, Biolegend, #345006, 4-1BB BV711, Biolegend, #309832, and eBioscience fixable viability dye 780, 1:1000, Thermo Fisher, #65-0865-18.

Surface staining was performed at 4°C for a minimum of 20 minutes, before cells were washed twice and acquired on a BD-LSR Fortessa. Data were then analysed in FlowJo (version 10).

### CRISPR-Cas9 mediated targeting of the SH2 domain gene family in primary CD8^+^ T-cells

Primary CD8^+^ T-cells were isolated from healthy donor leukocyte cones and transduced with lentiviruses encoding sgRNA molecules targeting all SH2 domain containing genes as previously described (**Supplementary Table 1**). Due to the size of this family, this analysis was split across 6 independent healthy CD8^+^ T-cell donors, where each donor was consistently assayed against an internal control sgRNA panel consisting of guides targeting: *LCK*, *LCP2*, *ITK*, *CD8* and *SHC4* together with an empty plasmid control. Following electroporation of Cas9 to induce perturbation of these genes, CD8^+^ T-cells were co-cultured with gp100 pulsed A375 melanoma cell lines, and the bispecific T-cell engager to a final concentration of 10^-10^M. Cytotoxic responses of CD8^+^ T-cells were assessed by measuring A375 melanoma growth, for a period of 48 hours. After 48 hours, CD8^+^ T-cells were harvested and stained with the following antibody cocktail at 4°C for 20 minutes to assess activation marker expression (CD8 PeCy7, 1:200, Biolegend #344712; CD25 AF647, 1:200, Biolegend #302618; 4-1BB BV711, 1:200, Biolegend #309832; eBioscience fixable viability dye 780, 1:1000, Thermo Fisher #65-0865-18).

### Donor-donor normalisations in primary CD8^+^ T-cell killing assays targeting SH2 domain family genes

In order to account for differing cytotoxic capacities between all CD8^+^ T-cell donors used to cover perturbation of the entire SH2 domain gene family, killing curves for each perturbation were first normalised to their corresponding internal negative control for each individual donor. This allows for the effect of targeting any given gene within the family to be compared with other genes in the family that were assayed on separate donors, given the size of the SH2 family. Next, to produce a single quantitative parameter for each gene, the area under the curve (AUC) was calculated for all normalised killing curves using the trapz function from the pracma package in R-studio (version 2024.09.1)(*48*). This analysis produced an individual value for targeting each gene within the family, which was used to determine whether this gene behaved as a negative or positive regulator of CD8^+^ T-cell responses (**Fig 3F**).

### Production of primary CD8^+^ 1G4-TCR^+^ T-cells and subsequent cytotoxicity assays

CD8^+^ T-cells isolated from healthy donor leukocyte cones were lentivirally transduced with plasmid encoding for the 1G4-TCR together with a LNGFR tag as previously described (*18*). Next, cells were then stained with TCR Vβ13.1 APC (Biolegend, #362408) and LNGFR FITC (Miltenyi Biotech, #130-113-423) at 1:200 and double positive events were purified by fluorescence-activated cell sorting. At the same time, PBMCs were isolated from three separate healthy donor leukocyte cones, and were subjected to an irradiation dose of 3000rad. Irradiated PBMCs were then centrifuged at 300g for 5 minutes, and resuspended at 2x10^6^/mL in complete RPMI-1640 medium. Equal volumes of PBMCs from each of the three donors was then combined, before supplementation of the mixture with 200U/mL IL-2, 1X PHA (500X stock, Invitrogen, #00-4977-03) and 50ng/mL OKT3 (Biolegend, #), to produce T-cell rapid expansion medium (REM). Approximately 1x10^5^ sorted LNGFR^+^ TCR Vβ13.1^+^ CD8^+^ T-cells was then added to 10mL REM and cultured for 5 days before counting and re-seeding to 10^6^ cells/mL in complete RPMI-1640 medium with 200U/mL IL-2. Cells were then expanded for a further 10-12 days, maintaining IL-2 concentrations at 200U/mL and cell densities at 1-2x10^6^/mL throughout. Resultant CD8^+^ T-cell cultures were checked for purity via flow cytometry and staining with CD8 PeCy7, TCR Vβ13.1 APC and LNGFR FITC. Functional reactivity was then assessed by co-culture with NY-ESO-1 expressing A375 cell lines prior to cryopreservation.

### IFNᵧ ELISAs

Supernatants were collected at 24 hours post CD8^+^ T-cell and A375 co-culture, and IFNᵧ secretion from CD8^+^ T-cells into the culture supernatant was assessed by ELISA as per manufacturer’s instructions (Biolegend, #430101). Supernatants were either used fresh in ELISAs, or were stored at -20°C prior to analysis but were never freeze-thawed.

### CyTOF sample preparation and data analysis

A375 cells were plated in 96-well flat-bottom plates at 5 × 10⁴ cells per well and pulsed with gp100 peptide (10µg/mL) for a minimum of 4 hours. After peptide loading, cells were washed once with PBS and either incubated with the bispecific T-cell engager at a final concentration of 10^-10^M or left untreated. Where applicable, bispecific binding was allowed to proceed for at least 30 minutes at 37 °C prior to initiation of the experiment.

Isolated CD8⁺ T cells were subsequently added to either gp100-pulsed bispecific-coated (stimulated) or uncoated (unstimulated) A375 cells and centrifuged (500 g, 1 min) to promote cell contact and initiate activation. At the end of the time course, cells were transferred to 96-well U-bottom plates, pelleted in ice-cold PBS (1800 rpm, 3 min), and fixed in 100 μl of 1.6% PFA. Fixation was carried out at 4 °C for a minimum of 20 minutes, followed by three washes with 200 μl PBS.

Mass cytometric analysis was performed as previously described. Briefly, dead cells were labelled with 0.25 mM ¹⁹⁴Cisplatin (Standard BioTools, Cat# 201194), after which cells were incubated overnight at 4 °C with thiol-reactive barcodes. Excess barcodes were quenched using 2 mM GSH and samples were pooled. Cells were then washed in cell staining buffer (CSB; Standard BioTools, Cat# 201068) and stained with extracellular rare-earth metal-conjugated antibodies for 30 minutes at room temperature. Cells were then permeabilised with 0.1% (v/v) Triton X-100/PBS and 50% methanol/PBS prior to staining with intracellular rare-earth metal-conjugated antibodies for 30 minutes at room temperature. Antibodies were cross-linked using 1.6% (v/v) FA/PBS for 10 minutes. Cells were subsequently incubated overnight at 4 °C with 125 nM ¹⁹¹Ir/¹⁹³Ir DNA intercalator, washed, resuspended in CAS+ supplemented with 2 mM EDTA, and analysed on a CyTOF XT at 200–400 events per second.

Signal intensities were measured by CyTOF at 1, 5, 10, 15, 30, 45, 60, and 120 minutes post-stimulation using a panel of 42 markers. Raw data were preprocessed in FlowJo and subsequently analysed using the CyGNAL python package (github.com/TAPE-Lab/CyGNAL(*14*)) with default settings.

### Analysis of published RNA expression and CRISPR dataset

To obtain the lists of signalome genes that were regulated during T cell activation, raw counts of bulk RNA sequencing dataset were downloaded from Gene Expression Omnibus (GEO). We filtered the data for CD8+ T cells except for GSE197067, where only pan-T cells were analyzed. Gene IDs were mapped to gene names by mygene package (Python). Data was prefiltered by subsetting for genes that have at least 10 counts in more than 20% of samples. Differential expression analysis was performed using DESeq2 (R)(*49*) using appropriate model design for study and p values were obtained with the wald test. Log fold change (LFC) was shrunk with the apeglm method to reduce inflation of lowly expressed genes. Differentially expressed genes (DEGs) were obtained by thresholding for adjusted p-value < 0.001, and LFC < 1.5. DEGs of different conditions (e.g. timepoints) of the same dataset were merged by their union and intersected with our signalome gene list. To obtain the lists of signalome genes that were identified as hits in pooled CRISPR screening, we downloaded the processed result files from each study, thresholded the hits following the description of each study and intersected with our signalome gene list.

## Author contribution

The experiments were conceived by S.S, E.P, S.J.D, and C.J.T, and the methodology developed by J.C and S.S. The experimental work was undertaken by J.C, S.S, C.T.G, E.B, M.K, A.M.S, M.T. V, P.S, J.X.L. Data analysis was performed by J.C, H.C, and S.S. The work was supervised by S.S, E.P, S.J.D, C.J.T, and funding was acquired by J.C, S.S, E.P, S.J.D, C.J.T. The original manuscript was written by J.C, S.S, and E.P and revised by H.C, M.K, A.M.S, C.J.T and S.J.D.

## Acknowledgements

This work was funded by MRC (grant MR/W025507/1 to E.P., S.S., C.T., and S.J.D.) and Wellcome grant to S.S (317519/Z/24/Z), and MSIF pump priming grant to J.C (BRDNB420).

