## Supplementary Figures for "Signalome-wide mapping of the NFκB pathway in T-cells reveals novel targets for immunotherapy"

Supplementary Fig 1: Generation of Cas9 expressing 1G4-TCR+ Jurkat NF-kB reporter cell line
Supplementary Fig 2: Recovery of core TCR signalling components in the screen.
Supplementary Fig 3: CyTOF-based single-cell analysis of T cell signalling states.
Supplementary Fig 4: Arrayed screening in primary T cells.
Supplementary Fig 5: Functional consequences of targeting *TRRAP* or *CTDSPL2* in primary CD8+ T-cells.

### Supplementary Tables

Supplementary Table 1: List of sgRNA sequences utilised in the signalome library
Supplementary Table 2: Hits identified from gene expression and CRISPR screen datasets
Supplementary Table 3: Arrayed screen data from both 9V and 6T stimulation condition.
Supplementary Table 3: RNA-seq dataset for TRRAP and CTDSPL2

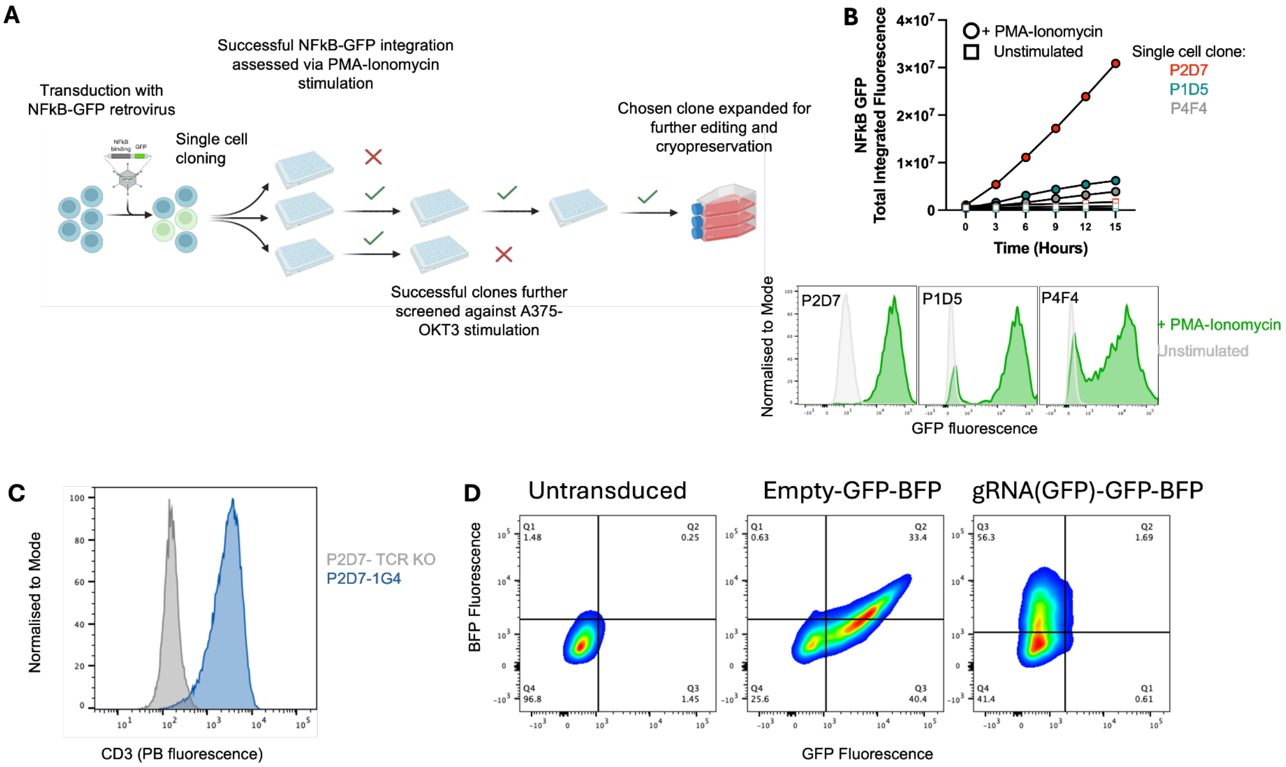


**Supplementary Fig. 1: Generation of Cas9 expressing 1G4-TCR⁺ Jurkat NF-kB reporter cell line. A:** Schematic overview of Jurkat NF-κB-GFP reporter generation. Jurkat cells were first transduced with retroviruses encoding the reporter construct and subsequently single-cell cloned. Successful integration of the reporter construct in proliferating clones was then assessed via screening of GFP expression in response to PMA-Ionomycin stimulation. GFP-positive clones were further expanded and subjected to secondary, TCR-specific stimulation using membrane-bound OKT3 presented on A375 cells. The clone demonstrating robust and specific activation was selected for further expansion. **B:** GFP expression following PMA/ionomycin stimulation in three representative clones. Clone P2D7 exhibited strong and uniform GFP induction (lower panel) and was selected for subsequent modification. **C:** Endogenous TCR in clone P2D7 was disrupted, and the 1G4-TCR was introduced to generate a 1G4-TCR$^{+}$ NF-κB reporter line. **D:** Cas9 was introduced into the 1G4-TCR reporter line via lentiviral transduction. Polyclonal Cas9-expressing populations were evaluated for genome editing efficiency using GFP-encoding lentiviral constructs carrying either a GFP-targeting gRNA (GFP) or no gRNA ("empty" control). Representative two-dimensional flow cytometry plots of BFP versus GFP fluorescence are shown. BFP marks cells expressing the gRNA construct. Efficient Cas9 activity is demonstrated by loss of GFP expression in cells transduced with the GFP-targeting gRNA.

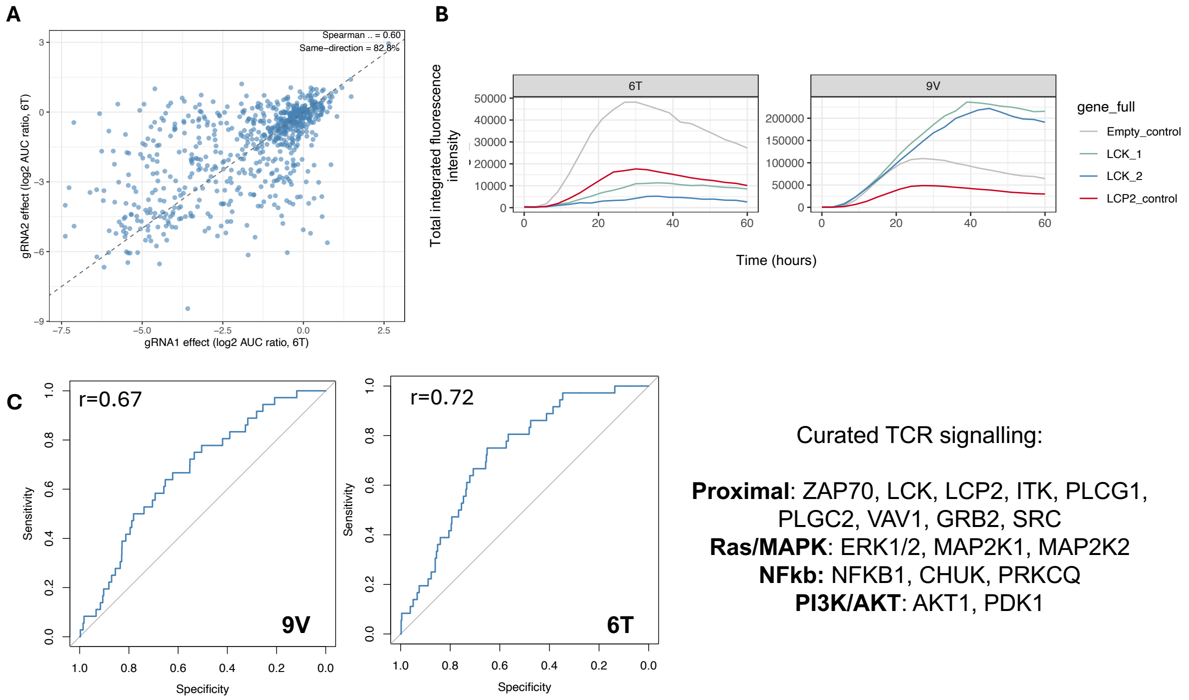


**Supplementary Fig.2: Recovery of core TCR signalling components in the screen. A:** Concordance between paired gRNAs under 6T peptide stimulation. Each point represents a gene; the dashed line indicates y = x. Spearman correlation coefficient (ρ) and percentage of same-direction effects are indicated. **B:** Representative time-course traces of integrated GFP fluorescence intensity for LCK gRNAs compared with empty control and the positive control (LCP2) following stimulation with 6T or 9V peptide. **C:** Receiver operating characteristic (ROC) curves showing the performance of the screen in identifying a curated set of TCR signalling genes under 6T (left) and 9V (right) stimulation conditions. The diagonal line indicates random classification.

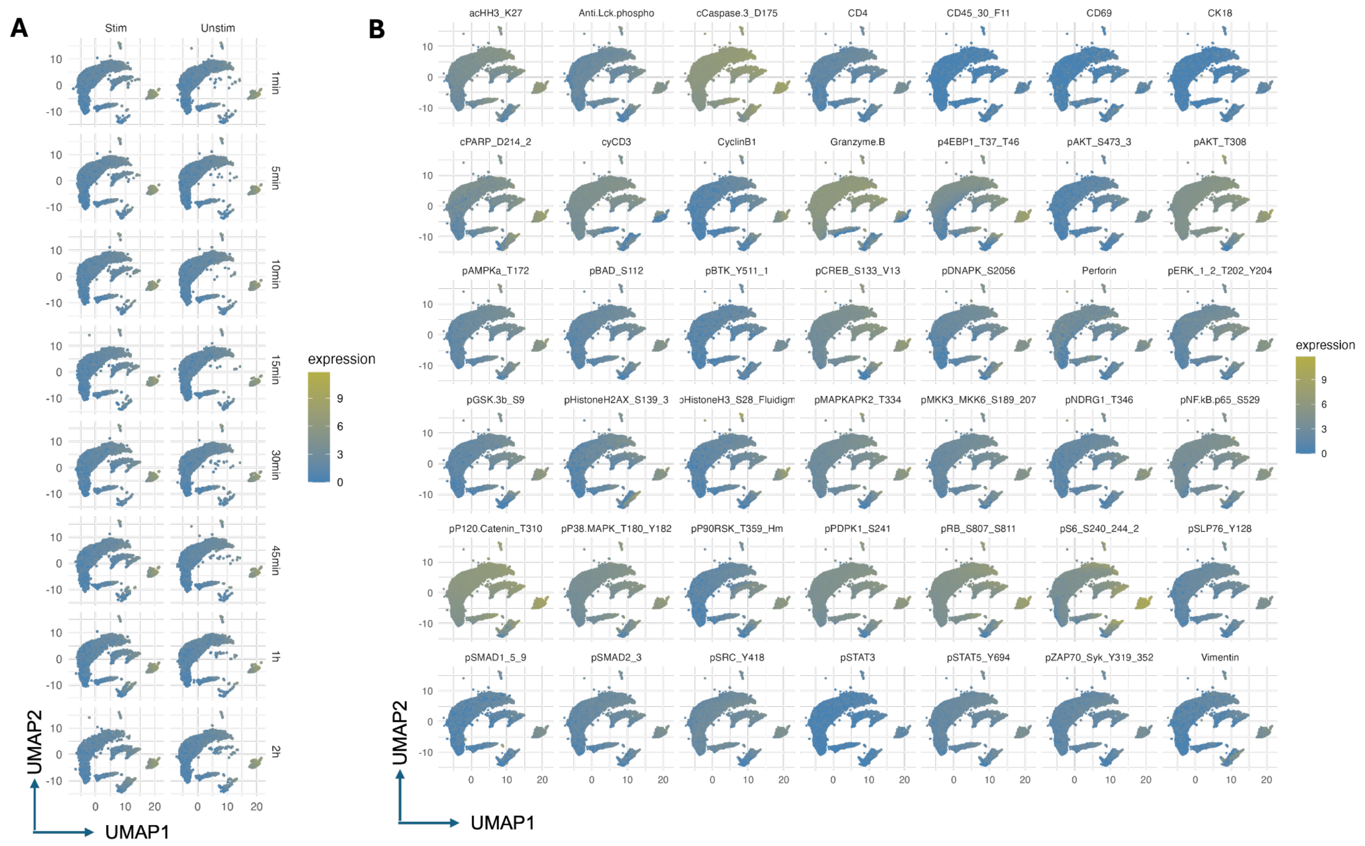


**Supplementary Fig.3 . CyTOF-based single-cell analysis of T cell signalling states.** Uniform Manifold Approximation and Projection (UMAP) visualization of CyTOF data from unstimulated (Unstim) and stimulated (Stim) T cells. Each point represents a single cell positioned according to high-dimensional marker expression profiles. UMAP was generated using all measured markers shown (arcsinh transformation, cofactor 5). Cells are coloured by arcsinh-transformed expression intensity of the indicated markers. A single global colour scale is applied across all panels to enable direct comparison of marker abundance between conditions and across markers.

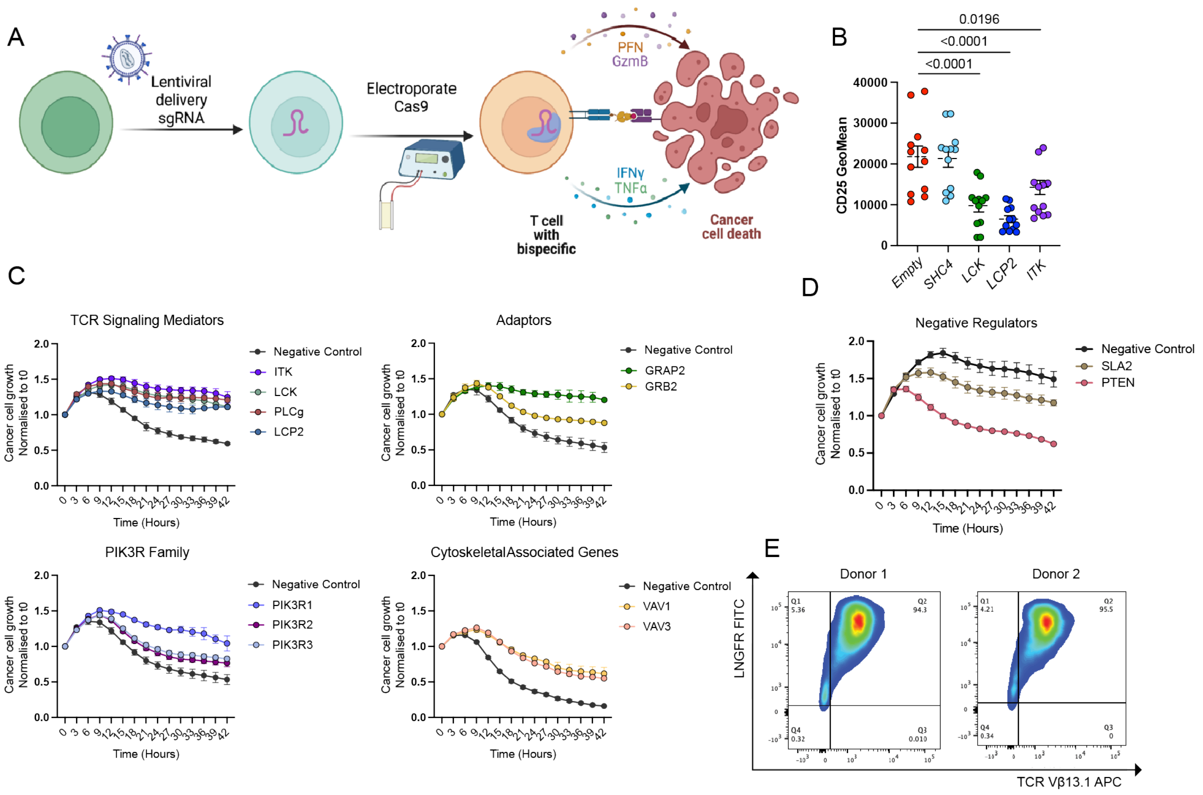


**Supplementary Fig. 4: Arrayed screening in primary T cells. A:** Schematic outlining the methodology used for CRISPR-Cas9 editing in primary CD8$^{+}$ T-cells and subsequent co-culture with A375 melanoma cells, and a bispecific T-cell engager for quantification of CD8$^{+}$ cytotoxic responses. **B:** Geometric mean of CD25 expression on CD8$^{+}$ T-cells 48 hours post-culture with A375 melanoma cells as described in A, with CRISPR perturbation of genes indicated. **C:** CD8$^{+}$ mediated killing of A375 melanoma cells using the pipeline illustrated in A, identifying positive regulators of T-cell activation. Data shown are representative curves of the entire SH2-gene family analysis shown in 2F. **D:** CD8$^{+}$ mediated killing of A375 melanoma cells as described in C, identifying negative regulators of T-cell activation. **E:** Representative gating and purity of primary CD8$^{+}$ 1G4-TCR$^{+}$ T-cells following transduction with 1G4/LNGFR encoding plasmids and subsequent expansion.

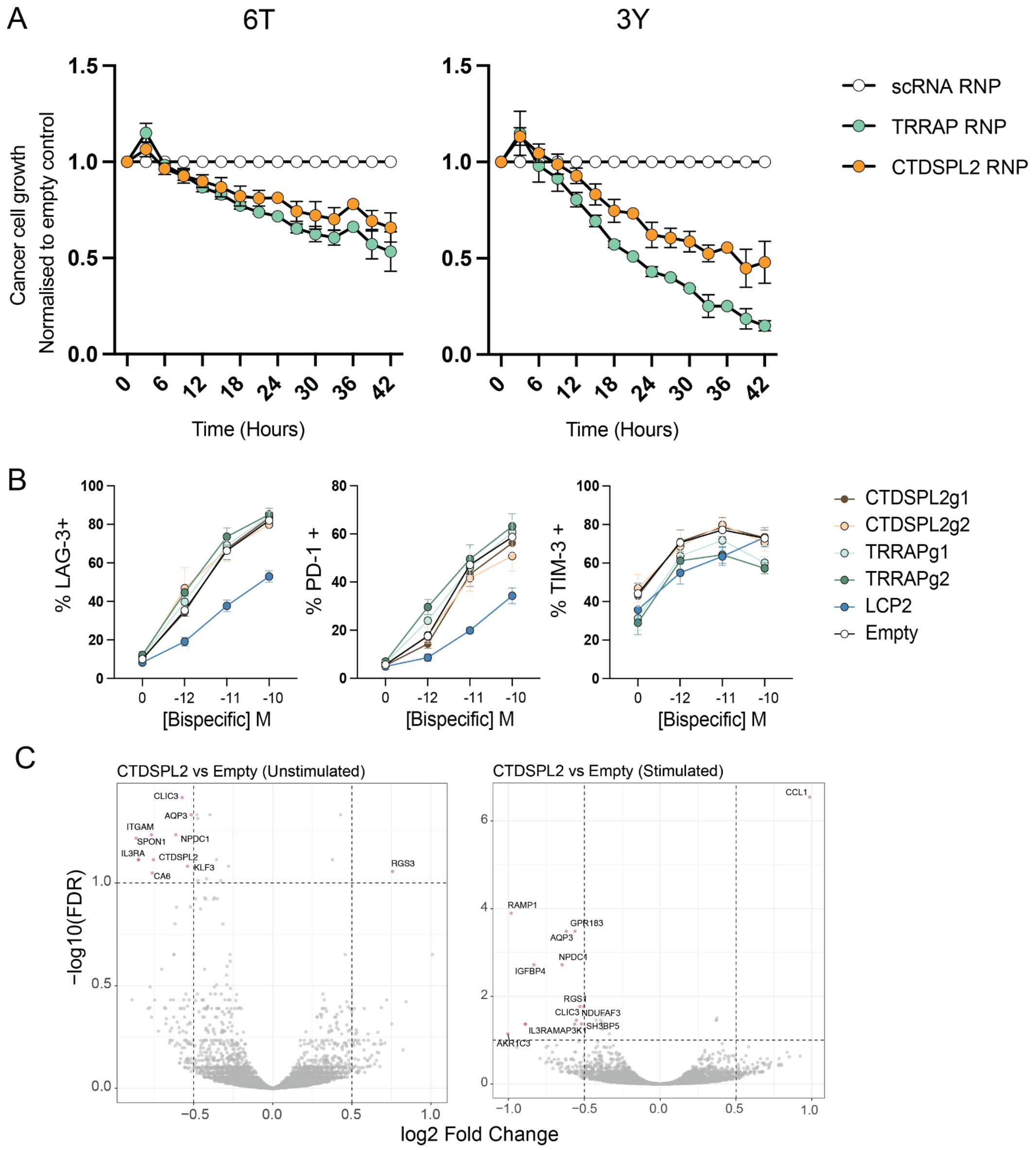


**Supplementary Fig. 5: Functional consequences of targeting *TRRAP* or *CTDSPL2* in primary $CD8^+$ T-cells. A:** Killing of A375 melanoma cells by 1G4-$TCR^+$ primary $CD8^+$ T-cells, with perturbation of *TRRAP* or *CTDSPL2* as indicated. Here, A375 melanoma cells were pulsed with either 6T or 3Y NY-ESO-1 peptides (100µg/mL), and resulting killing curves were normalised to the scRNA RNP control. Data shown from 2 independent donors. **B:** Surface expression of LAG-3, PD-1 and TIM-3 on primary $CD8^+$ T-cells with CRISPR-Cas9 targeting of indicated genes. $CD8^+$ T-cells were co-cultured with gp100 pulsed A375 melanoma cells and a bispecific T-cell engager to concentrations indicated, and surface inhibitory receptor expression was assessed 24 hours post co-culture. Data shown from 3 independent $CD8^+$ T-cell donors, and representative of a minimum of 2 independent experiments. **C:** Volcano plots depicting differentially expressed genes in CTDSPL2 targeted primary T cells (2 donors and 2 guides each) compared to empty controls (matched 2 donors) depicting minimal changes in transcriptomics either in unstimulated (left panel) or post 3 hour stimulated condition (right panel).
